# Control of the signaling of RAS proteins by modulating their palmitoylation

**DOI:** 10.1101/2024.06.27.600952

**Authors:** Jiakai Zhu, Ruiying Guo, Qi Hu

## Abstract

Small GTPases, such as the RAS family proteins, are key regulators of cell proliferation and oncogenesis. S-palmitoylation, a reversible post-translational modification catalyzed by ZDHHC enzymes, plays an important role in the regulation of many small GTPases by modulating their subcellular localization. However, development of chemical tools to modulate S-palmitoylation remains challenging due to the redundancy and poor druggability of ZDHHC enzymes. Here, we developed an approach to control the signaling of RAS proteins by modulating their palmitoylation through fusing acyl-protein thioesterase 1 (APT1) to their N-termini. Using *in vitro* and cell assays, we showed that the subcellular localization and signaling of NRAS in the fusion protein can be reversibly controlled by an APT1 inhibitor, ML348. This approach enables the observation of temporal changes in RAS protein localization, providing a useful tool for studying RAS signaling. Furthermore, similar approaches may be used to control the signaling of other small GTPases.

## Introduction

Small GTPases are molecular switches that regulate various cellular processes, such as cell growth and vesicular trafficking^1^. They switch between an active GTP-bound state and an inactive GDP-bound state, with the switch rates tightly regulated by guanine nucleotide exchange factors (GEFs) and GTPase-activating proteins (GAPs)^1^. In addition, S-palmitoylation, a reversible post-translational modification, also plays a crucial role in modulating the activity of many small GTPases^2-8^. S-palmitoylation is the covalent attachment of a long-chain acyl group, usually a palmitoyl group, to specific cysteine residues within the proteins via a thioester bond^9,10^. This reaction is catalyzed by a class of enzymes known as palmitoyltransferases, also named ZDHHC enzymes because of the conserved zinc-finger and the Asp-His-His-Cys sequence in the catalytic center of each enzyme^9,10^. We refer to S-palmitoylation as palmitoylation in our study for short. The palmitoylated small GTPase, with the help of the hydrophobic palmitoyl group, can localize to membranes such as the plasma membrane and Golgi membrane and mediate cell signaling. The palmitoylation can be reversed by another class of enzymes called depalmitoylases^11^. The ZDHHC enzymes and the depalmitoylases coordinate to control the palmitoylation levels and thus regulate the activities of small GTPases.

Among small GTPases, the RAS family proteins, including HRAS, NRAS, KRAS4a and KRAS4b, are of particular interest due to their important roles in cell proliferation and cancer treatment^12^. RAS proteins each consist of a highly conserved G-domain, which is responsible for the intrinsic GTPase activity, and a C-terminal hypervariable region (HVR)^12^. The CAAX motif at the end of the HVR of each RAS protein can be modified by prenylation at the cysteine residues, and then processed by RAS converting CAAX endopeptidase 1 (RCE1) to remove the AAX sequence, followed by carboxyl terminal methylation catalyzed by isoprenylcysteine carboxyl methyltransferase (ICMT)^12^. Additionally, cysteine residues adjacent to the CAAX motif in HRAS, NRAS, and KRAS4a, but not KRAS4b, can be modified by palmitoylation^12^. Palmitoylation and depalmitoylation are essential for the correct subcellular localization of HRAS and NRAS^2^. Mutation of the palmitoylation site in the G12D oncogenic mutant of NRAS to serine disrupted its oncogenic activity^13^.

Although the importance of palmitoylation in the regulation of cell signaling has been widely recognized, there is currently a lack of chemical tools for the precise modulation of protein palmitoylation. The difficulty lies in the redundancy and poor druggability of the ZDHHC enzymes. There are 23 human ZDHHC enzymes that have been identified^10^. Palmitoylation of RAS proteins is known to be catalyzed by the ZDHHC9/GCP16 complex^14^. We recently found that the ZDHHC14/GCP16 and the ZDHHC18/GCP16 complexes can also catalyze the palmitoylation of HRAS and NRAS^15^. So far, only a few broad-spectrum ZDHHC inhibitors have been reported^16^. An alternative strategy is to target the substrates of ZDHHC enzymes. A successful case is the development of covalent inhibitors of the stimulator of interferon genes protein (STING) to block its palmitoylation^17^. But for most palmitoylated proteins, such as small GTPases, palmitoylation occurs at flexible regions, making it quite difficult to identify small molecule inhibitors.

In this study, we developed an approach to reversibly control the signaling of RAS proteins by modulating their palmitoylation. We fused acyl-protein thioesterase 1 (APT1), which has been reported to be a depalmitoylase of RAS proteins^18^, to the N-terminus of NRAS to create a fusion protein. We showed that in cells, palmitoylation of NRAS in this fusion protein and its ability to activate the downstream ERK1/2 were blocked by the fused APT1, but can be turned on by adding an APT1 inhibitor ML348 (ref.^19^) and turned off again by the removal of ML348.

We also demonstrated that the APT1 fusion protein strategy can be used to regulate the palmitoylation of two other RAS proteins: KRAS4a and HRAS.

## Results

### NRAS can be depalmitoylated by fusing the depalmitoylase APT1 to its N-terminus

We first checked the subcellular localization of NRAS carrying the oncogenic mutation Q61R. A FLAG-tagged green fluorescent protein (GFP) was added to the N-terminus of NRAS(Q61R) to facilitate monitoring of the subcellular localization. The fusion protein, called GN, was localized to the plasma membrane and Golgi when stably expressed in HEK293T cells (Figure 1a, b). Golgi localization was indicated by co-localization with the Golgi marker GalT-blue fluorescent protein (BFP) that was co-expressed with the fusion protein GN. In contrast, when residue Cys181 in NRAS, which is the palmitoylation site of NRAS, was mutated to serine, the fusion protein GN^C181S^ was mainly localized in the cytosol (Figure 1a, b), indicating that the subcellular localization of NRAS is regulated by palmitoylation at Cys181. These results are consistent with findings from a previous study^2^.

**Figure 1.**
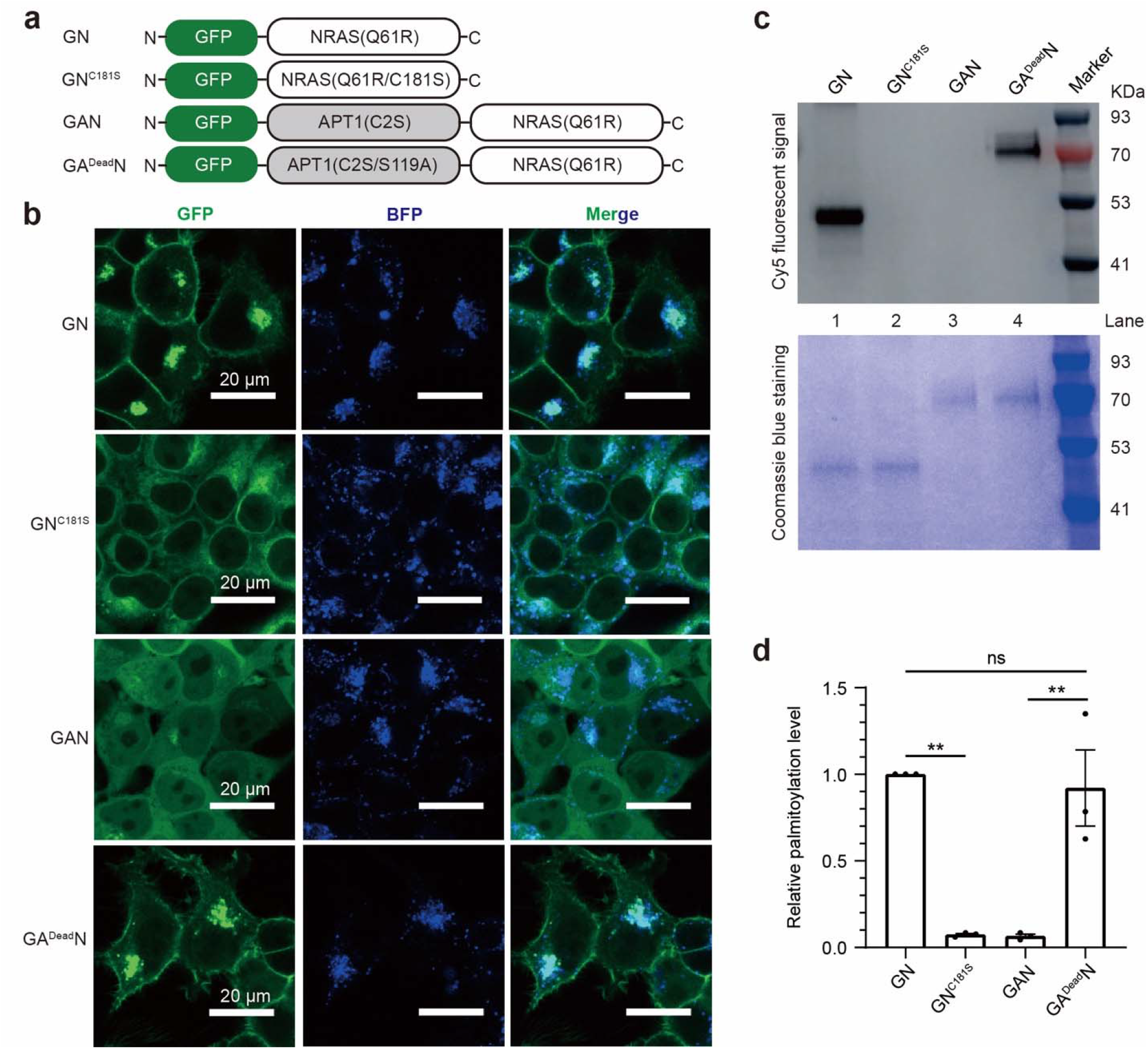
Fusing APT1 to NRAS disrupts the membrane localization of NRAS. **a**, The linear diagrams of the NRAS fusion proteins. **b**, Subcellular localization of the NRAS fusion proteins stably co-expressed with the Golgi marker GalT-BFP in HEK293T cells. The GFP signals indicate the localization of the NRAS fusion proteins, while the BFP signals indicate the localization of the Golgi marker GalT-BFP. **c, d**, The palmitoylation levels of the NRAS fusion proteins were assessed by supplementing the cell culture medium with 17-Octadecynoic Acid (17-ODYA), which is used as a substitute for palmitic acid. Then the fusion proteins modified by 17-ODYA were labeled with a fluorescent dye Cy5 via click reaction. The data in (**c**) are the results of a representative experiment out of three independent experiments. The data in (**d**), which represent the mean ± SD of three independent measurements, were analyzed using the one-way ANOVA in Prism to calculate the two-tailed P values: **, P < 0.01; ns, P ≥ 0.05.

Next, we tested whether bringing the human depalmitoylase APT1 into proximity with NRAS can promote the depalmitoylation of NRAS. We fused APT1 to the N-terminus of NRAS(Q61R). APT1 has been reported to be regulated by palmitoylation at Cys2, and upon palmitoylation, APT1 localizes to the cell membrane^20^. In the fusion protein, we mutated this Cys2 to serine to block the palmitoylation of APT1. Additionally, we added a GFP to the N-terminus of APT1 in the fusion protein. The resulting fusion protein was called GAN (Figure 1a). A control protein, GA^Dead^N, was constructed by mutating the catalytic residue Ser119 in APT1 in the fusion protein GAN to alanine (Figure 1a). GAN was mainly localized in the cytosol when stably expressed in HEK293T cells; in contrast, GA^Dead^N was localized to the plasma membrane and Golgi, similarly to the fusion protein GN (Figure 1b). These results demonstrate that APT1 fused to the N-terminus of NRAS can disrupt the membrane localization of NRAS, and this effect relies on the catalytic activity of APT1.

We also checked the palmitoylation levels of these NRAS fusion proteins when they were stably expressed in HEK293T cells using a click assay (Figure 1c, d). GN and GA^Dead^N showed similar palmitoylation levels, while the palmitoylation of GN^C181S^ and GAN was not detectable. These results are consistent with the subcellular localization data, suggesting that APT1 regulates the subcellular localization of NRAS by catalyzing the depalmitoylation of NRAS.

### NRAS in the fusion protein can be activated by an APT1 inhibitor

We speculate that inhibiting the catalytic activity of APT1 using small molecule inhibitors may restore the palmitoylation level of NRAS in the fusion protein. To test this idea, we first evaluated the palmitoylation of the fusion protein GAN using NBD-palmitoyl-CoA as a fluorescent analog of palmitoyl-CoA in an *in vitro* reconstituted assay^15^. GAN and GA^Dead^N were overexpressed in *E. coli* and purified to homogeneity. The purified GAN or GA^Dead^N was incubated with ML348 — a selective inhibitor of human APT1 (ref.^19^) or the same volume of DMSO, and then mixed with the purified ZDHHC9/GCP16 complex. Palmitoylation was initiated by adding NBD-palmitoyl-CoA. After 1 hour, palmitoylation of GAN and GA^Dead^N was detected using fluorescent imaging (Figure S1). The palmitoylation level of GAN was almost undetectable in the absence of ML348 (Figure S1a, lanes 1 and 2), but significantly increased in the presence of ML348 (Figure S1a, lanes 3 and 4). In contrast, the palmitoylation level of GA^Dead^N was significantly increased in the absence of ML348 after 1 hour, and adding ML348 did not further increase the palmitoylation level (Figure S1a, lanes 5-8; Figure S1b). These results indicate that inhibition of APT1 can upregulate the palmitoylation of NRAS in the fusion protein GAN.

Next, we evaluated the effect of the APT1 inhibitor ML348 on the subcellular localization of GAN in HEK293T cells. Upon treating the HEK293T cells stably expressing GAN with 10 µM of ML348, GAN re-localized from the cytosol to the Golgi in a time-dependent manner (Figure 2a, b). In contrast to the accumulation of GAN on the Golgi, GAN on the plasma membrane did not increase after ML348 treatment (Figure 2a, b). Thirty minutes after adding ML348, the subcellular localization of the fusion protein GAN became similar to that of GN and GA^Dead^N (Figures 1b and 2a).

**Figure 2.**
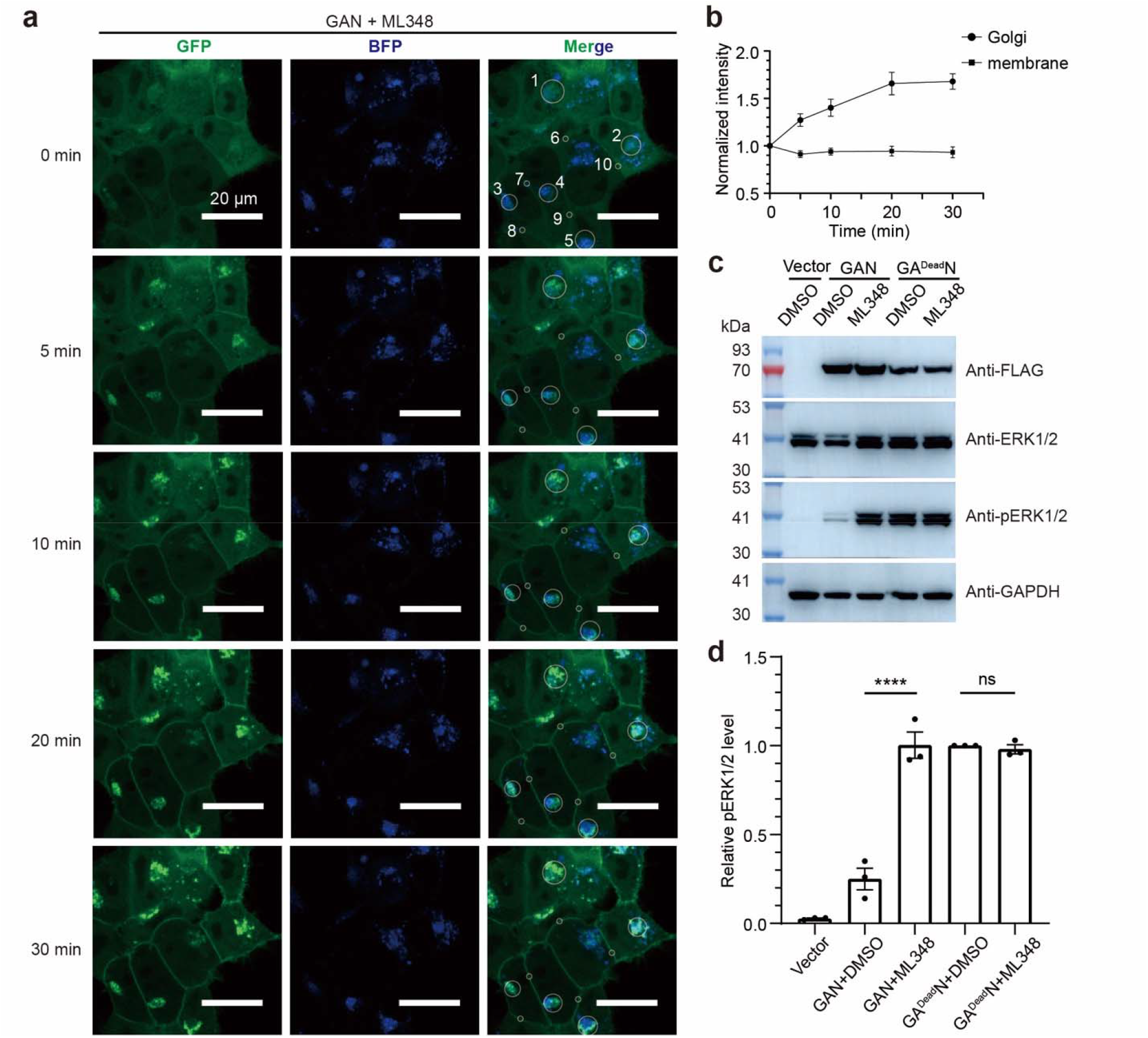
APT1 inhibitor ML348 activates the palmitoylation of NRAS fusion protein GAN. **a**, Changes in the subcellular localization of the NRAS fusion protein GAN stably expressed in HEK293T cells upon treatment with the 10 µM of APT1 inhibitor ML348. The white circles numbered 1 to 5 indicate the location of the Golgi, while those numbered 6 to 10 indicate the location of the plasma membrane. **b**, Quantification of the changes in GFP signals on the Golgi and plasma membrane after adding 10 µM of ML348. The data in (**b**) represent the mean ± SEM of signals indicated by white circles in (**a**). **c** and **d**, Activation of ERK1/2 in HEK293T cells by stably expressing the NRAS fusion proteins GAN and GA^Dead^N, followed by treating the cells with ML348. The expression of GAN and GA^Dead^N was examined by the anti-FLAG antibody. The data in (**c**) are the results of a representative experiment out of three independent experiments. The data in (**d**) represent the mean ± SD of three independent measurements and were analyzed using the one-way ANOVA in Prism to calculate the two-tailed P values: ****, P < 0.0001; ns, P ≥ 0.05.

We also checked whether the re-localization of GAN induced by ML348 could activate the downstream RAF-MEK-ERK pathway. Indeed, the phosphorylation levels of the extracellular signal-regulated kinases 1 and 2 (ERK1/2) in HEK293T cells stably expressing GAN were significantly increased upon 30 minutes of ML348 treatment (Figure 2c, d). In contrast, the phosphorylation of ERK1/2 in HEK293T cells stably expressing GA^Dead^N remained high regardless of ML348 treatment (Figure 2c, d). We next analyzed the kinetics of ERK1/2 phosphorylation after treating the HEK293T cells with ML348. In contrast to HEK293T cells that did not express GAN, the phosphorylation level of ERK1/2 in HEK293T cells stably expressing GAN had a jump between 10 and 20 minutes, and then remained at a high level until 720 minutes (Figure S2).

Lin and Conibear reported that ABHD17 proteins, rather than APT1 and APT2, are depalmitoylases that catalyze NRAS depalmitoylation^21^. To test the effect of ABHD17 proteins on the subcellular localization of the NRAS fusion protein GAN, we pretreated the HEK293T cells with ABD957, a covalent inhibitor of the ABHD17 proteins^22^, for 4 hours, then treated the cells with ML348 (Figure S3a, b). GAN was slightly accumulated on the plasma membrane, with its intensity on the plasma membrane increasing by approximately 20% after 30 minutes of treatment with ML348, suggesting that GAN on the plasma membrane can be depalmitoylated by ABHD17 depalmitoylases. However, more obvious accumulation of GAN was observed on the Golgi (Figure S3a, b), similarly to that in the absence of ABD957 (Figure 2a, b).

To test the effect of ABHD17 inhibition on the signaling activity of GAN, HEK293T cells carrying the empty vector or stably expressing GAN were pretreated with DMSO, 0.5 µM, or 10 µM of ABD957, followed by treatment with ML348 or DMSO (Figure S3c, d). In the absence of GAN expression, no ERK1/2 phosphorylation was detected, regardless of treating the cells with ABD957 and ML348 (Figure S3c). In the presence of GAN, 10 µM of ABD957 alone slightly increased the phosphorylation level of ERK1/2; however, treating the cells with both ABD957 and ML348 did not further increase the ERK1/2 phosphorylation level compared to treatment with only ML348.

The above results demonstrate the feasibility of modulating the NRAS signaling pathway by fusing NRAS with the depalmitoylase APT1 and treating the fusion protein with selective APT1 inhibitors. These results also indicate that the increased ERK1/2 phosphorylation level in HEK293T cells stably expressing GAN upon treatment with ML348 might mainly come from GAN localized to the Golgi, rather than to the plasma membrane.

### NRAS without fusing APT1 is insensitive to the APT1 inhibitor ML348

When applying the fusion protein GAN in the study of NRAS palmitoylation, one concern is that ML348 also inhibits endogenous APT1, and thus its effect on the palmitoylation of proteins beyond GAN may not be ignored. In addition, the depalmitoylase activity of APT1 in GAN may not be limited to GAN when expressed in human cells. To address these concerns, we fused FLAG-tagged mCherry to the N-terminus of NRAS(Q61R) and transiently co-expressed this fusion protein, called mCN, with GAN and the Golgi marker GalT-BFP in HEK293T cells. We then checked whether the subcellular localization of mCherry-NRAS(Q61R) is affected by ML348 treatment (Figure 3).

**Figure 3.**
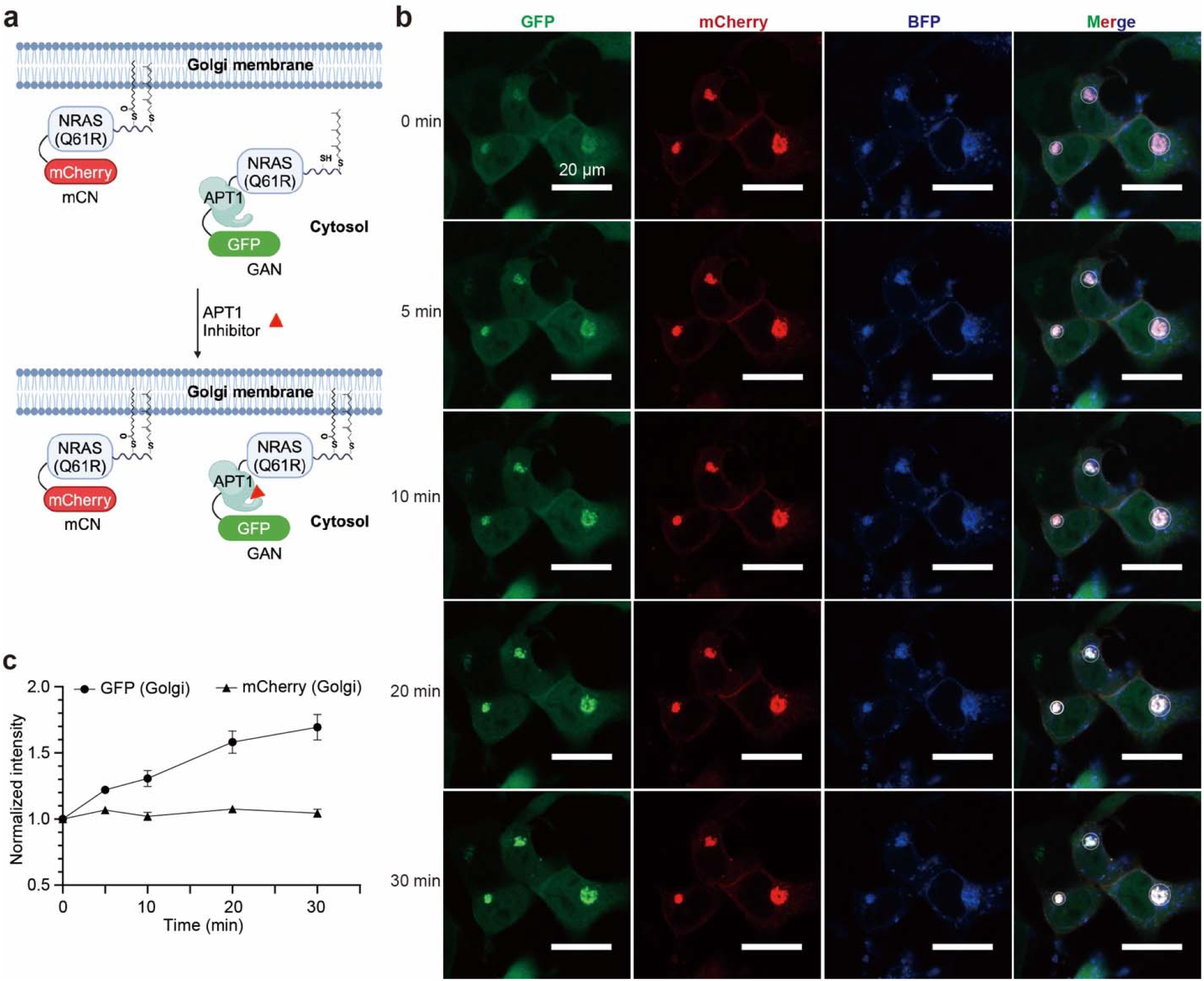
ML348 treatment does not affect the localization of NRAS without fusing APT1. **a**, Illustration of the regulation of NRAS localization by ML348. **b**, Changes in the subcellular localization of the NRAS fusion proteins GAN and FLAG-mCherry-NRAS(Q61R), also called mCN, transiently expressed in HEK293T cells upon adding 10 µM of ML348. **c**, Quantification of the changes in the GFP and mCherry signals on the Golgi in the HEK293T cells after adding 10 µM of ML348. The data in (**c**) represent the mean ± SEM of signals in three cells indicated by white circles in (**b**).

GAN was mainly localized in the cytosol at the beginning, and after adding ML348, it accumulated on the Golgi (Figure 3b, c). This result is consistent with what was observed when GAN was stably expressed in HEK293T cells (Figure 2a, b). In contrast, mCN was mainly localized to the plasma membrane and Golgi (Figure 3b). The intensity of the mCherry signal on the Golgi was not obviously changed upon ML348 treatment (Figure 3c). Similar results were observed when GAN and mCN were transiently co-expressed in Hela cells (Figure S4). These results suggest that NRAS without fusing APT1 is insensitive to the APT1 inhibitor ML348.

### The APT1 inhibitor ML348 can reversibly modulate NRAS localization

To test whether the ML348-induced re-localization of the RAS fusion protein GAN is reversible, we removed ML348 by changing the cell culture medium after treating the cells with ML348 for 30 minutes (Figure 4a). Upon treating the HEK293T cells, stably expressing GAN and the Golgi marker GalT-BFP, with 1 µM of ML348, GAN quickly accumulated on the Golgi (Figure 4b, d), similar to Figure 2a. After ML348 was removed, GAN gradually redistributed to the cytosol. (Figure 4c, d).

**Figure 4.**
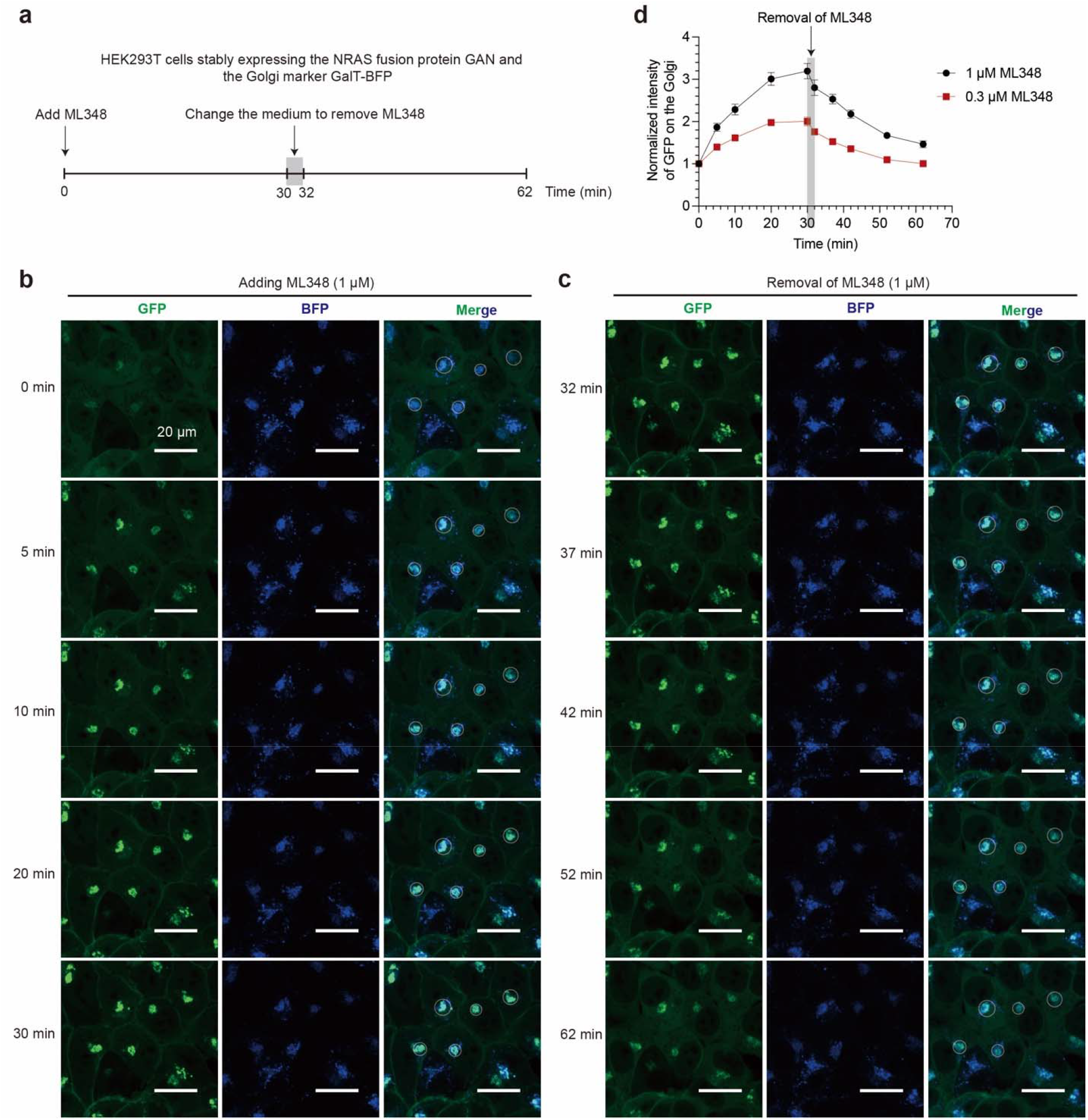
ML348 induced re-localization of NRAS fusion protein GAN is reversible. **a**, Illustration of the cell imaging process after adding or removing ML348. **b**,**c**, Changes in the subcellular localization of the NRAS fusion protein GAN stably expressed in HEK293T cells after adding 1 µM of ML348 (**b**), and subsequently removing ML348 (**c**). **d**, Quantification of the changes in the GFP signals on the Golgi after adding and subsequently removing ML348. The data in (**d**) represent the mean ± SEM of signals in five cells indicated by white circles in (**b**), (**c**) and Figure S5.

To demonstrate the dose dependence of the effect of ML348, we also treated the HEK293T cells with 0.3 µM of ML348 (Figure S5a). GAN also accumulated rapidly on the Golgi, but the accumulation was weaker compared to that in the presence of 1 µM of ML348 (Figures 4d and S5b). Removal of the 0.3 µM of ML348 from the culture medium also led to redistribution of GAN to the cytosol (Figures 4d and S5c).

### KRAS4a and HRAS have different sensitivities to APT1 fused to their N-termini

We also tested the effectiveness of the APT1 fusion protein strategy on KRAS4a and HRAS, the other two RAS proteins known to be regulated by palmitoylation^12^. The G12C mutant of KRAS4a, with a GFP-tagged APT1 fused to its N-terminus, was mainly localized in the cytosol when stably expressed in HEK293T cells. It was re-localized to the plasma membrane and Golgi upon ML348 treatment (Figures 5a and S6a). However, the G12C mutant of HRAS, even with a GFP-tagged APT1 fused to its N-terminus, was localized on the plasma membrane and Golgi regardless of ML348 treatment (Figures 5b and S6b). This could be explained by the dual palmitoylation sites in HRAS. In contrast to NRAS and KRAS4a, HRAS has two palmitoylation sites: C181 and C184 (ref.^12^). It has been reported that neither the C181S nor the C184S mutation alone disrupted the membrane localization of HRAS^2^.

**Figure 5.**
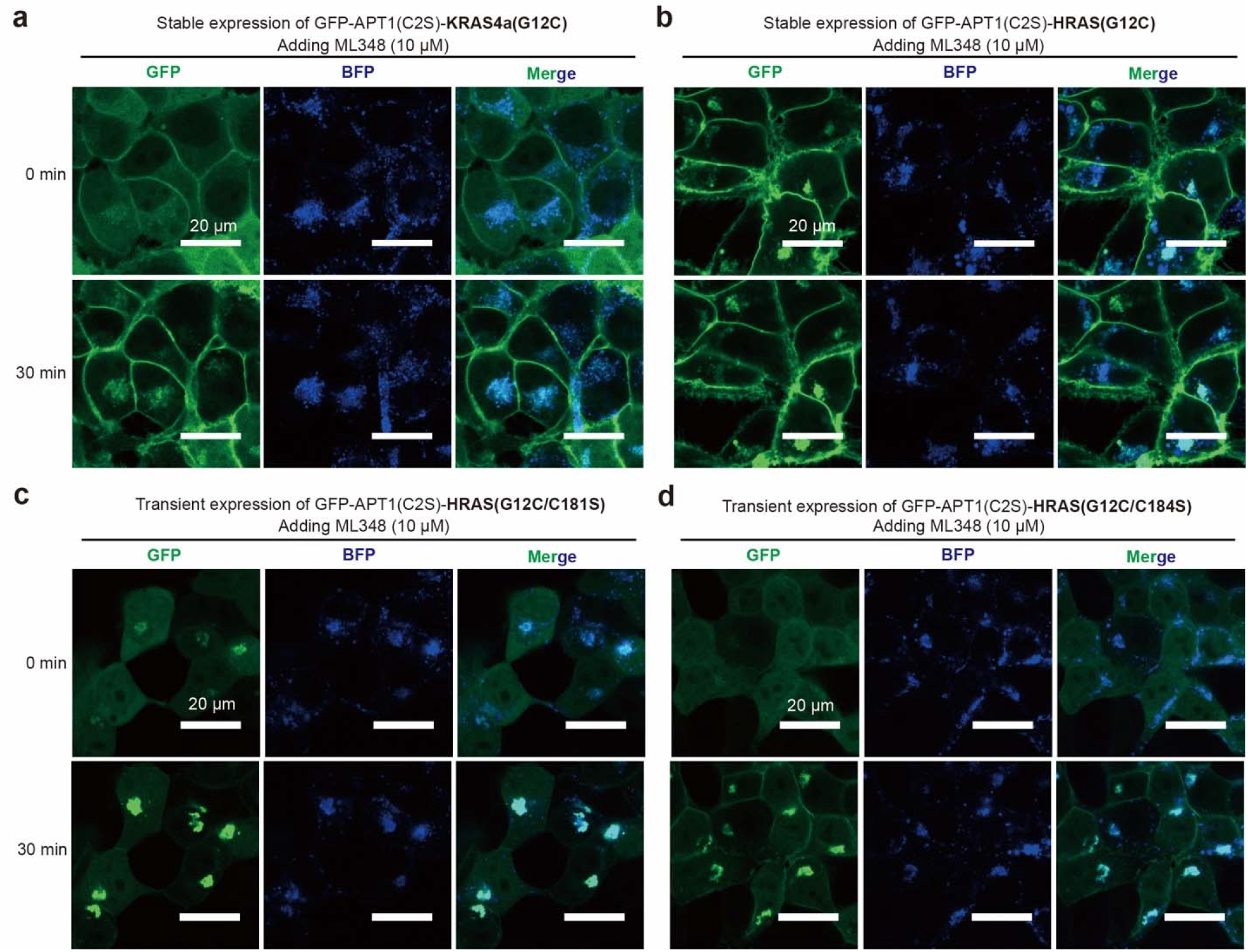
Fusing APT1 can also promote the depalmitoylation of KRAS4a and HRAS. **a,b**, KRAS4a(G12C) (**a**) and HRAS(G12C) (**b**) each with a GFP-tagged APT1(C2S) fused to the N-terminus were stably co-expressed with the Golgi marker GalT-BFP in HEK293T cells. **c, d**, The HRAS(G12C/C181S) (**c**) and the HRAS(G12C/C181S) (**d**) double mutants each with a GFP-tagged APT1(C2S) fused to the N-terminus were transiently co-expressed with the Golgi marker GalT-BFP in HEK293T cells. After adding 10 µM of ML348, the cells were imaged using ZEISS LSM 980 Confocal Microscope. The cell images taken at 5 min, 10 min, and 20 min after adding ML348 can be found in Figures S6 and S7.

Next, we evaluated the effect of fusing APT1 on the subcellular localization of HRAS(G12C) carrying either the C181S or the C184S mutation (Figures 5c, d and S7). With a GFP-tagged APT1 fused to their N-termini, both the C181S and the C184S mutants were almost completely distributed in the cytosol when they were transiently expressed in HEK293T cells. After treating the cells with 10 µM of ML348, the G12C/C181S mutant re-localized to the Golgi, while the G12C/C184S mutant re-localized to both the Golgi and plasma membrane. The effects of the C181S and C184S mutations on the subcellular localization of HRAS observed in our study are consistent with previous findings^2^. These findings suggest that the fused APT1 can catalyze the depalmitoylation of HRAS, but it is not sufficient to simultaneously maintain both C181 and C184 at their depalmitoylated states. There might be other depalmitoylases in cells that contribute to the depalmitoylation of HRAS.

## Discussion

The importance of palmitoylation and depalmitoylation of RAS proteins has been well recognized^12^, however, how the palmitoylation and depalmitoylation processes are regulated is largely unclear. In this study, we have shown that bringing the depalmitoylase APT1 into proximity with RAS proteins promotes the depalmitoylation of them, thus disrupting their membrane localization. Creation of the APT1-RAS fusion proteins enables reversible manipulation of the localization of RAS proteins using a selective APT1 inhibitor, ML348. This approach allows us to observe the temporal changes in RAS localization. Thus, it would be useful for studying the dynamics of RAS palmitoylation and depalmitoylation and for screening factors that regulate these dynamics.

Our data also allowed us to uncover a surprising signaling role of Golgi-localized NRAS. In our system, chemical rescue of NRAS palmitoylation increased its Golgi localization without changing the amount on the plasma membrane. Meanwhile, we observed a ∼4x increase in pERK levels (Figure 2). A recent report suggests that Golgi-localized RAS confers resistance to small molecule KRAS inhibitors^23^. Further investigation is clearly merited to understand the cell signaling mediated by the Golgi-localized NRAS.

Although the ZDHHC enzymes and depalmitoylases have been well known for their regulatory roles in numerous cellular processes^10^, no selective ZDHHC small molecule regulators have been reported, and only a few small molecule inhibitors of depalmitoylases have been identified^16,22,24^. Development of methods for quickly evaluating the cellular activities of ZDHHCs or depalmitoylases would facilitate the discovery of small molecule regulators. A few fluorescent probes have been developed to study the S-depalmitoylation activity in live cells^25-27^, however, these probes cannot discriminate the S-depalmitoylation activity of different depalmitoylases. Our APT1-RAS fusion protein approach presents a high-throughput method for screening regulators of APT1 and ZDHHC enzymes that catalyze RAS palmitoylation. Compounds that effectively inhibit or enhance the enzymatic activity of APT1 or ZDHHC enzymes could be rapidly assessed based on their impact on the membrane localization of the GAN fusion protein.

Considering the conserved structures of small GTPases, our approach may also be used to regulate the palmitoylation of other small GTPases. Furthermore, we demonstrated that fusing an enzyme with its substrates provides a convenient way to selectively and reversibly regulate the function of a specific substrate using an inhibitor of the enzyme.

Lastly, our study suggests that development of bifunctional molecules to recruit depalmitoylases to NRAS is a feasible strategy to inhibit NRAS signaling and would be promising for the treatment of cancers carrying NRAS activating mutations.

## Acknowledgments

We would like to thank Dr. Ziyang Zhang at University of California, Berkeley for valuable comments. We would like to thank Guicun Fang at the Westlake University Microscopy Core Facility for advice and assistance in light microscopy sample preparation and data collection. This work was supported by Westlake Laboratory of Life Sciences and Biomedicine, Westlake Education Foundation, and “Pioneer” and “Leading Goose” R&D Program of Zhejiang (2023C03109, 2024SSYS0036).

## Author Contributions Statement

Q.H. conceived and supervised the project; J.Z. designed and performed the assays; R.G. performed the assays; all authors contributed to data analysis; Q.H. and J.Z. wrote the manuscript.

## Data availability

All other data are available in the manuscript or the supplementary materials.

