## Supplementary Material for "Control of the signaling of RAS proteins by modulating their palmitoylation"

**This PDF file includes:**

Materials and Methods

Figures S1 to S5

**Materials and Methods:**

Cell culture

The HEK293T cells were cultured in high-glucose DMEM (Hyclone, Cat# C11995500BT) supplemented with 10% FBS (Gibco, Cat# 10270-106) and 1% penicillin-streptomycin solution (Hyclone, Cat# SV30010). After recovery from liquid nitrogen, the cells were cultured in a humidified incubator with 5% $CO_2$ at 37 °C for at least two generations before they were used in the assays. The HEK293F cells were cultured in SMM293-TII medium (Sino Biological, Cat# M293TII) supplemented with 1% penicillin-streptomycin solution at 37 °C under 5% $CO_2$ in a Multitron-Pro shaker (Infors, 5% $CO_2$, 110 rpm).

Genes and cloning

The UniProt numbers of the proteins used in this study include APT1 (UniProt No.: O75608), NRAS (UniProt No.: P01111), KRAS4a (UniProt No.: P01116-1), HRAS (UniProt No.: P01112), GalT (UniProt No.: P15291, 1-82 aa), BFP (FPbase ID: ZO7NN), GFP (FPbase ID: R9NL8, UniProt No.: C5MKY7).

For protein purification, the plasmids of the ZDHHC9/GCP16 and NRAS were constructed following a protocol described in a previous study of our group[1]; the genes encoding GFP-APT1 (C2S)-NRAS(Q61R) and GFP-APT1(C2S/S119A)-NRAS(Q61R) were cloned into a modified pET15b vector to express proteins with an N-terminal 6x His tag.

For transient expression in HEK293T and Hela cells, the genes encoding FLAG-GFP-APT1(C2S)-NRAS(Q61R), FLAG-mCherry-NRAS(Q61R), FLAG-GFP-APT1(C2S)-HRAS(G12C/C181S), FLAG-GFP-APT1(C2S)-HRAS(G12C/C184S), and GalT-BFP were cloned into a pcDNA3.1 vector.

For stable expression in HEK293T cells, the genes encoding the fusion proteins, including FLAG-GFP-NRAS(Q61R), FLAG-GFP-NRAS(Q61R/C181S), FLAG-GFP-APT1(C2S)-NRAS(Q61R), FLAG-GFP-APT1(C2S/S119A)-NRAS(Q61R), FLAG-GFP-APT1(C2S)-KRAS4a(G12C), and FLAG-GFP-APT1(C2S)-HRAS(G12C), were cloned into the lentiviral vector pLVX-Puro, while the gene encoding GalT-BFP (synthesized by Genewiz) was cloned into the lentiviral vector pLVX-neo.

The linker sequence between the FLAG tag and GFP (or mCherry), between GFP (or mCherry) and NRAS or APT1, and that between APT1 and NRAS, KRAS4a or HRAS, are all Gly-Gly-Gly-Ser.

Generation of stably transduced HEK293T cells

To generate HEK293T cells stable expressing target proteins, after the cells in 10-cm dish were cultured to 60-80% confluency, a mixture of 9 µg pLVX-DNA, 4 ug pVSVG and 6 ug pMD8.9 dissolved in 1 mL of Opti-MEM (no FBS; no P/S) was mixed with 19 µL of Linear Polyethylenimine 25,000 (Polysciences, Cat# 23966) dissolved in 1 mL of Opti-MEM (no FBS; no P/S) for 20 min at room temperature. The mixture was added to the cultured 293T cells, and the medium was exchanged after transfection at 8 h. The cells were cultured for 48 h after transfection, then the supernatant was harvested and filtered through a 0.45-µm syringe filter. The filtrate containing the lentivirus was used immediately or stored at -80 °C. 1 mL of the lentivirus of each target gene was added to 1x $10^6$ per well HEK293T cells (6-well plate) in the presence of 10 µg/mL of polybrene (Merck, Cat# TR-1003-G). After 24 h, the medium was exchanged and the cells were cultured for additional 24 h, and then 2 µg/mL of puromycin was added for selection. For cells stably transfected with both the target gene and the Golgi marker GalT-BFP, flow cytometry was used to obtain cells expressing both the target protein and GalT-BFP after two rounds of transduction.

Western blot

The cells from 6-well plates were centrifugated at 600 g for 5 min at 4 °C and washed 3 times with cold PBS, then lysed by adding 100 μL of the RIPA lysis buffer (50 mM Tris-HCl pH 7.4, 150 mM NaCl, 1% Triton X-100, 0.5% sodium deoxycholate, 0.1% SDS) supplemented with the Complete Protease Inhibitor Cocktail (Roche, Cat# 04693116001) on ice for 20 min. Following centrifugation at 22,000 g at 4 °C for 10 min, the concentration of the total protein in the supernatant was determined by the BCA assay (Thermo Fisher Scientific, Cat# 23225 and Cat# 23227). Each sample containing equal amount of total protein was mixed with a 2x loading buffer and boiled at 95 °C for 5 min, and then were subjected to 4-20% SDS-PAGE gels (GenScript, Cat# M00657) and transferred to a 0.2-μm PVDF membrane (Merck Millipore). After blocking with 5 % milk in TBST buffer, the PVDF membranes were incubated with antibodies of target proteins for further WB analysis. The primary antibodies used in this study include Anti-FLAG (Sigma, Cat# F1804), Anti-His (GenScript, Cat# A00186-100), Anti-GADPH (Immunoway, Cat# YM3029) Anti-MAPK (ERK1/2) (CST, Cat# 4695S), and Anti-phospho-MAPK (pERK1/2) (Thr202/Tyr204) (CST, Cat# 4370S). The Anti-phospho-MAPK (pERK1/2) (Thr202/Tyr204) antibody detects dually phosphorylated ERK1 (at Thr202 and Tyr204) and ERK2 (at Thr185 and Tyr187), and mono-phosphorylated ERK1 (at Thr202) and ERK2 (at Thr185). The secondary antibodies include HRP-conjugated anti-mouse antibody (Merck Millipore, Cat# AP127P) and HRP-conjugated anti-rabbit antibody (Merck Millipore, Cat# AP156P).

For the kinetic analysis of ERK phosphorylation in HEK293T cells, the cells carrying the empty vector or stably expressing GAN were grown in 3.5-mm dishes and treated with 10 μM of ML348. At designated time points (0 min, 5 min, 10 min, 20 min, 30 min, 1 h, 3 h, 6 h, and 12 h), the cells were washed with cold PBS. After washing, the cells were centrifuged at 800 g for 2 minutes at 4 °C, then immediately frozen in liquid nitrogen. For cell lysis and Western blotting, the protocols were the same as described above.

For the ABD957 treatment assay, HEK293T cells carrying the empty vector or stably expressing GAN were cultured in 6-well plates. After pretreating the cells with 0.5 μM or 10 μM of ABD957 or the same volume of DMSO for 4 hours, 10 μM of ML348 was added to the medium. 30 minutes after adding ML348, the cells were centrifuged at 800 g for 2 minutes at 4°C and promptly frozen in liquid nitrogen. The subsequent steps for cell lysis and Western blotting were the same as described above.

Protein expression and purification

The ZDHHC9/GCP16 complex and NRAS were overexpressed and purified following the protocol describe in a previous study of our group[1]. GFP-APT1(C2S)-NRAS(Q61R) and GFP-APT1(C2S/S119A)-NRAS(Q61R) each in a modified pET15b vector were transformed into *E. coli* BL21(DE3) cells. The cells were cultured in LB medium supplemented with 0.1 mg/mL ampicillin at 37 °C until $OD_{600}$ reached 0.8-1, then cooled to 18 °C followed by addition of 200 μM β-D-thiogalactopyranoside (IPTG). The cells were cultured at 18 °C overnight, then harvested by centrifugation at 4,000 g for 10 min at 4 °C, resuspended in a lysis buffer (25 mM Tris-HCl pH7.4, 150 mM NaCl) and lysed by ultrasonication. The cell lysates were centrifuged at 22,000 g at 4 °C for 1 h (BECKMAN Avanti JXN-26 Centrifuge), then the supernatant was incubated with

TALON Metal Affinity Resin (TaKaRa, Cat# 635504) and the target protein was purified following the manufacturer's protocol. The eluate was supplemented with 5 mM of Dithiothreitol (DTT), loaded into a Source-15Q column (GE Healthcare) and eluted by a linear gradient from 100% buffer A (25 mM Tris-HCl pH 7.4) to 60% Buffer B (25 mM Tris-HCl pH 7.4, 1 M NaCl). The peak fractions were pooled, supplemented with 5 mM DTT, and further purified by a Superdex-200 increase 10/300 GL column (GE Healthcare) with a running buffer containing 25 mM HEPES pH 7.4 and 150 mM NaCl. The peak fractions were pooled and used for biochemical assays.

NBD-palmitoyl-CoA assay

The NBD-palmitoyl-CoA assay was carried out following a protocol described previously with slight modifications[1]. Specifically, 2 μM of the purified NRAS or NRAS fusion proteins was firstly incubated with 10 μM of ML348 or the same volume of DMSO in a reaction buffer (200 mM $NaH_2PO_4$, pH 7.2) for 30 min at 37 °C, then mixed with 0.2 μM of the ZDHHC9/GCP16 complex. After adding 1 μM of NBD-palmitoyl-CoA, half of the reaction mixture was immediately quenched by adding 2x SDS-PAGE loading buffer and boiling at 95 °C for 5 min (the sample was marked as the 0 h sample), while the other half was incubated at room temperature for 1 h and then quenched (the sample was marked as the 1 h sample). Then samples were subjected to 4-20% SDS-PAGE gels for further analysis. The gels were first imaged using Amersham Imager 680 (Cytiva) with a Cy2 filter (Excitation: 460 nm), and then stained by Coomassie blue.

Click assay

HEK293T cells cultured in 6-well plates were metabolically labeled with 20 μM of 17-ODYA in DMEM (0.2% BSA, no palmitic acid, no FBS) for 3 h. The cells were then collected by centrifugation at 600 g for 5 min and washed 3 times with DPBS. Each sample was lysed on ice with 100 μL of RIPA lysis buffer supplemented with Complete Protease Inhibitor Cocktail and 20 μM palmostatin B (Merk, 508738). The cell lysates were centrifugated at 13,000g for 10 min at 4 °C. The supernatants were collected, and the total protein concentration of each sample was measured using BCA assay. For each sample, an equal amount of total protein was incubated with 30 μL of anti-FLAG beads at 4 °C for 2 h. After centrifugation at 4,000 g for 8 s at room temperature, the the beads were washed with RIPA lysis buffer (1 mL x 3). The beads were then incubated with a click mix (1 mM TCEP, 0.1 mM TBTA, 1 mM $CuSO_4$, 0.1 mM Cy5-azide) for 1 h at room temperature. After the reaction, the beads were washed with RIPA lysis buffer (1 mL x 3), and then were boiled with 2x SDS-PAGE loading buffer at 95 °C for 5 min and subjected to 4-20% SDS-PAGE gels. The gels were first imaged using Amersham Imager 680 (Cytiva) with a Cy5 filter (Excitation: 630 nm), and then stained by Coomassie blue.

Confocal imaging

HEK293T or Hela cells were cultured in high-glucose DMEM supplemented with 10% FBS and 1% penicillin-streptomycin solution in confocal culture dishes (Biosharp, BS-15-GJM) until the confluency reached about 60%. Before imaging, the medium was exchanged to DMEM (1.9 mL per well, no FBS), and the cells were cultured for 1 h. After adding 100 μL of a stock of ML348 (200 μM, 20 μM, or 6 μM) in DMEM (no FBS) to each well, the cells were imaged using Axio ObserverZ1/7 microscope and Plan-Apochromat 40x/0.95 Korr M27 objective with an image size of 212.13 × 212.13 $\mu m^2$ on ZEISS LSM 980 Confocal Microscope. The cells were photographed for 30 min with a frequency of one image per min. For the subcellular localization analysis, the Golgi localization was designated by the spline contour tool in the ZEISS ZEN 3.8 software. The fluorescence intensity of GFP or mCherry at each time point was normalized to the intensity at 0 min to show the changes in the subcellular localization of the target proteins.

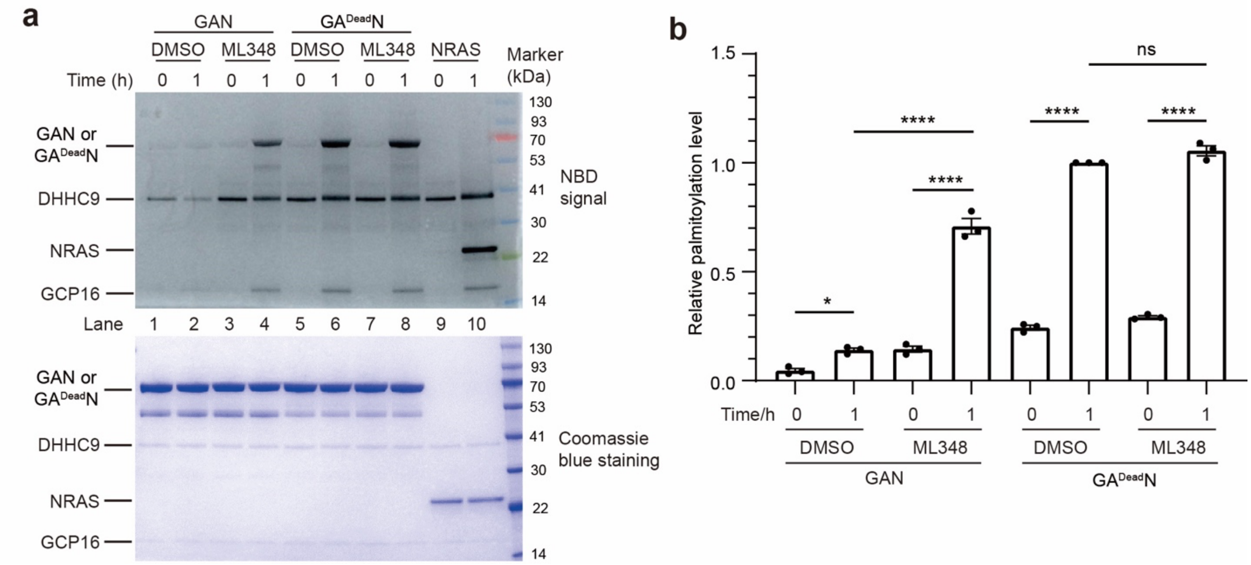


**Figure S1. APT1 fused to NRAS can abolish the palmitoylation of NRAS**. **a**, Palmitoylation of the purified NRAS fusion proteins catalyzed by the ZDHHC9/GCP16 complex in the absence or presence of 10 μM of the APT1 inhibitor ML348. NBD-palmitoyl-CoA was used as a substitute of palmitoyl-CoA. After the reaction, the samples were separated by SDS-PAGE and the gel was imaged using fluorescence imaging to quantify the NBD-palmitoylation level (top). The same gel was stained with Coomassie blue to show the loading amount of each protein (bottom). The data in (**a**) are the results of a representative experiment out of three independent experiments. **b**, Quantification of the NBD-palmitoylation levels of GAN and GA$^{Dead}$N. The data in (**b**) represent the mean ± SD of three independent measurements and were analyzed using the one-way ANOVA in Prism to calculate the two-tailed P values: ****, P < 0.0001; *, P < 0.05; ns, P ⩾ 0.05.

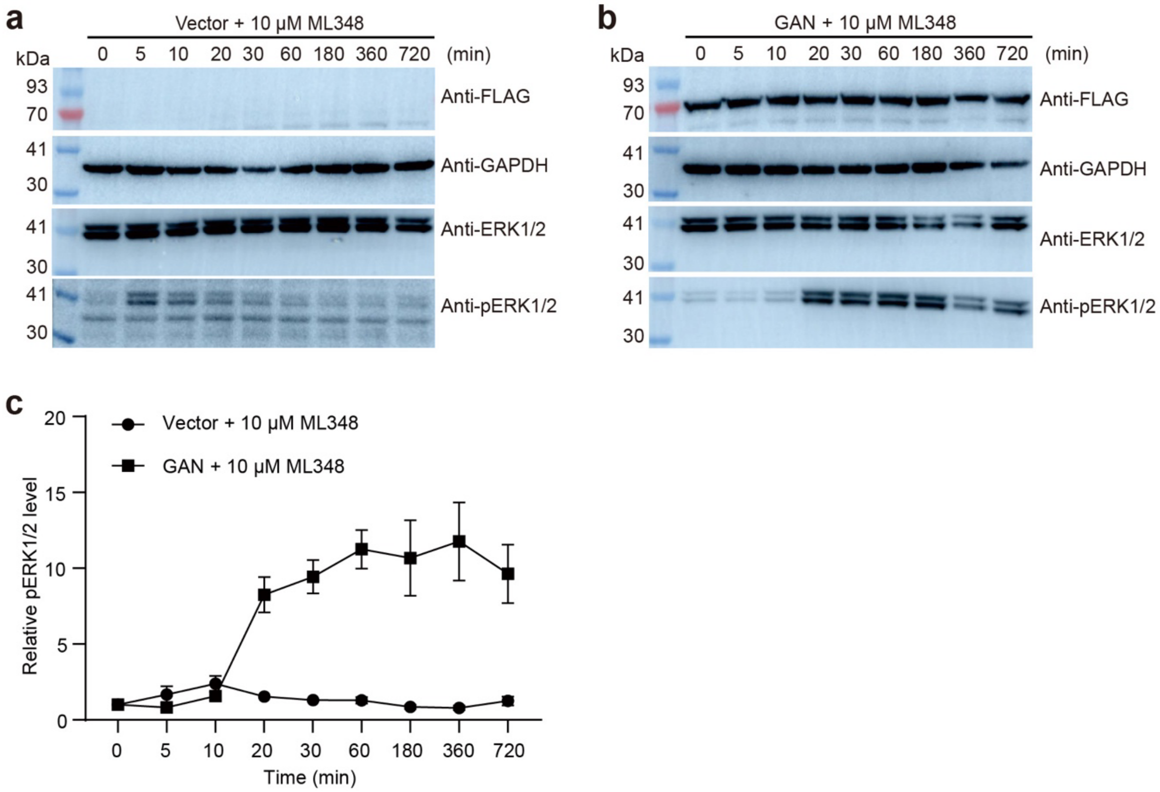


**Figure S2. Kinetics of ERK1/2 phosphorylation. a** and **b**, HEK293T cells carrying the empty vector (a) or stably expressing GAN (b) were treated with 10 μM of the APT1 inhibitor ML348 and then the phosphorylation of ERK1/2 was monitored using Western Blot. The expression of GAN was examined using the anti-FLAG antibody. The data in (**a**) and **(b**) are the results of a representative experiment out of three independent experiments. **c**. The phosphorylation levels of ERK1/2 in (**a**) and **(b**) were quantified and normalized to the levels at 0 minute. The data in (**c**) represent the mean ± SEM of three independent measurements.

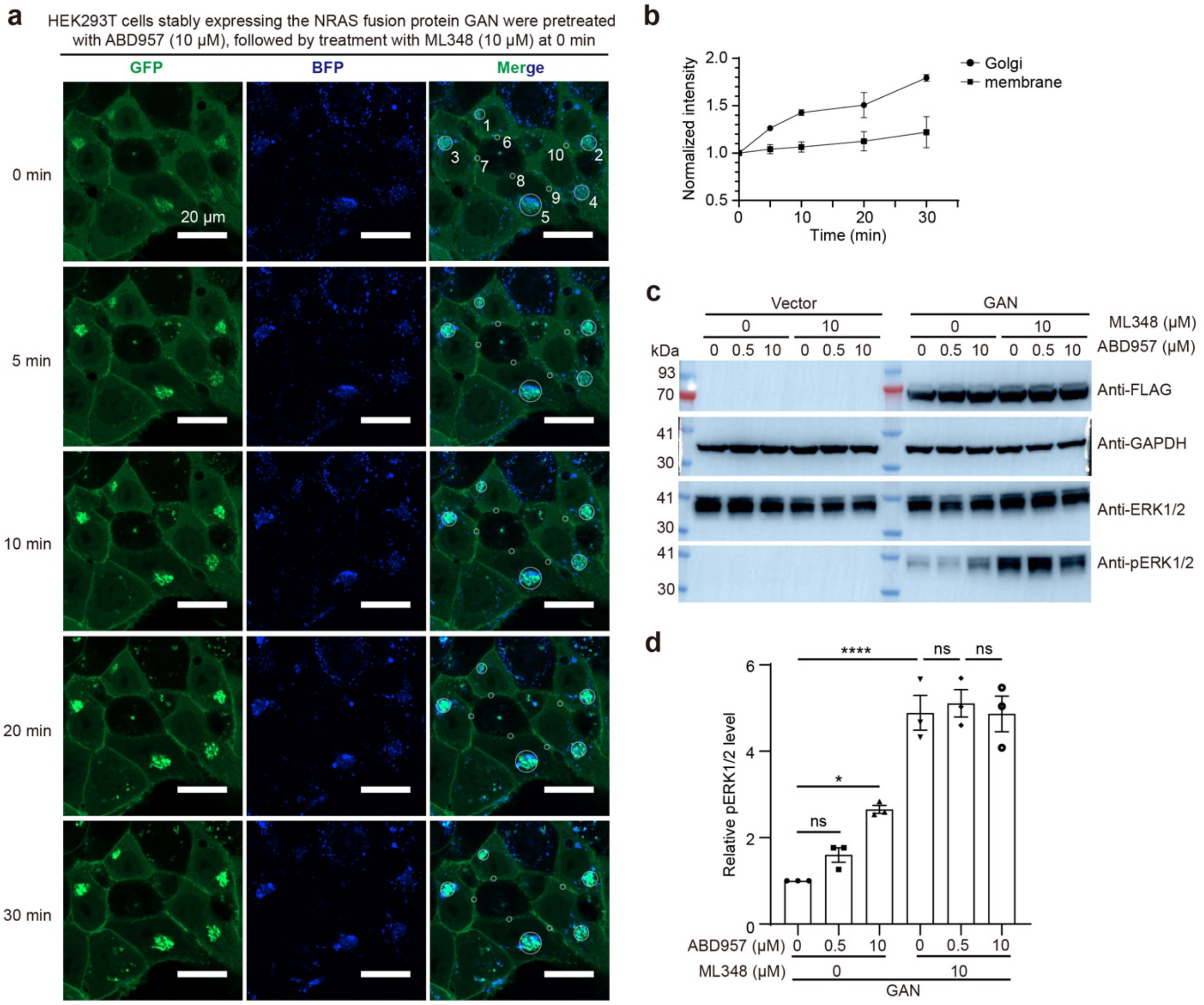


**Figure S3. Effects of the ABHD17 inhibitor ABD957 and the APT1 inhibitor ML348 on the subcellular localization and signaling activity of GAN.** **a**, Changes in the subcellular localization of the NRAS fusion protein GAN stably expressed in HEK293T cells upon pre-treating the cells with ABD957 for 4 hours, followed by treating the cells with ML348. The white circles numbered 1 to 5 indicate the location of the Golgi, while those numbered 6 to 10 indicate the location of the plasma membrane. **b**, Quantification of the changes in GFP signals on the Golgi and plasma membrane after adding 10 μM of ML348. The data in (**b**) represent the mean ± SEM of signals indicated by white circles in (**a**). **c**, HEK293T cells carrying the empty vector or stably expressing the NRAS fusion protein GAN were pretreated with different concentrations of

ABD957, and then treated with ML348 or the same volume of DMSO for 30 minutes. Then the phosphorylation of ERK1/2 was monitored using Western Blot. The expression of GAN was examined using the anti-FLAG antibody. The data in (**c**) are the results of a representative experiment out of three independent experiments. **d**, The phosphorylation levels of ERK1/2 in (**c**) were quantified and normalized to the levels in the absence of ABD957. The data in (**d**) represent the mean ± SEM of three independent measurements and were analyzed using the one-way ANOVA in Prism to calculate the two-tailed P values: ****, $P < 0.0001$; *, $P < 0.05$; ns, $P \geq 0.05$.

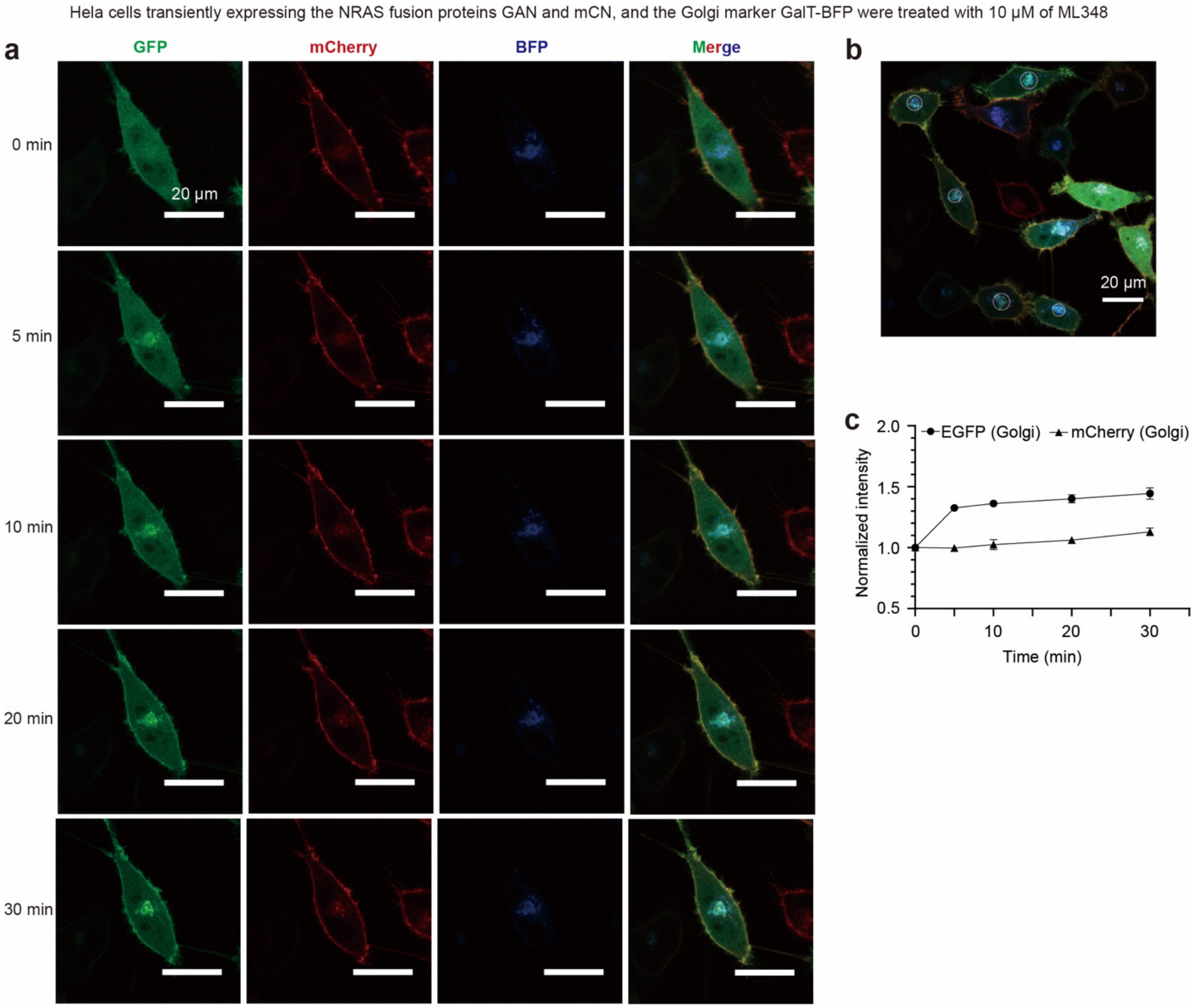


**Figure S4. ML348 treatment does not affect the localization of NRAS without fusing APT1 in Hela cells. a**, Changes in the subcellular localization of the NRAS fusion proteins GAN and FLAG-mCherry-NRAS(Q61R), also called mCN, transiently expressed in Hela cells upon adding 10 μM of ML348. **b**, The GFP and mCherry signals on the Golgi in five cells were recorded and used in the analysis in (**c**). **c**, Quantification of the changes in the GFP and mCherry signals on the Golgi in the Hela cells after adding 10 μM of ML348. The data in (**c**) represent the mean ± SEM of signals in five cells indicated by white circles in (**b**).

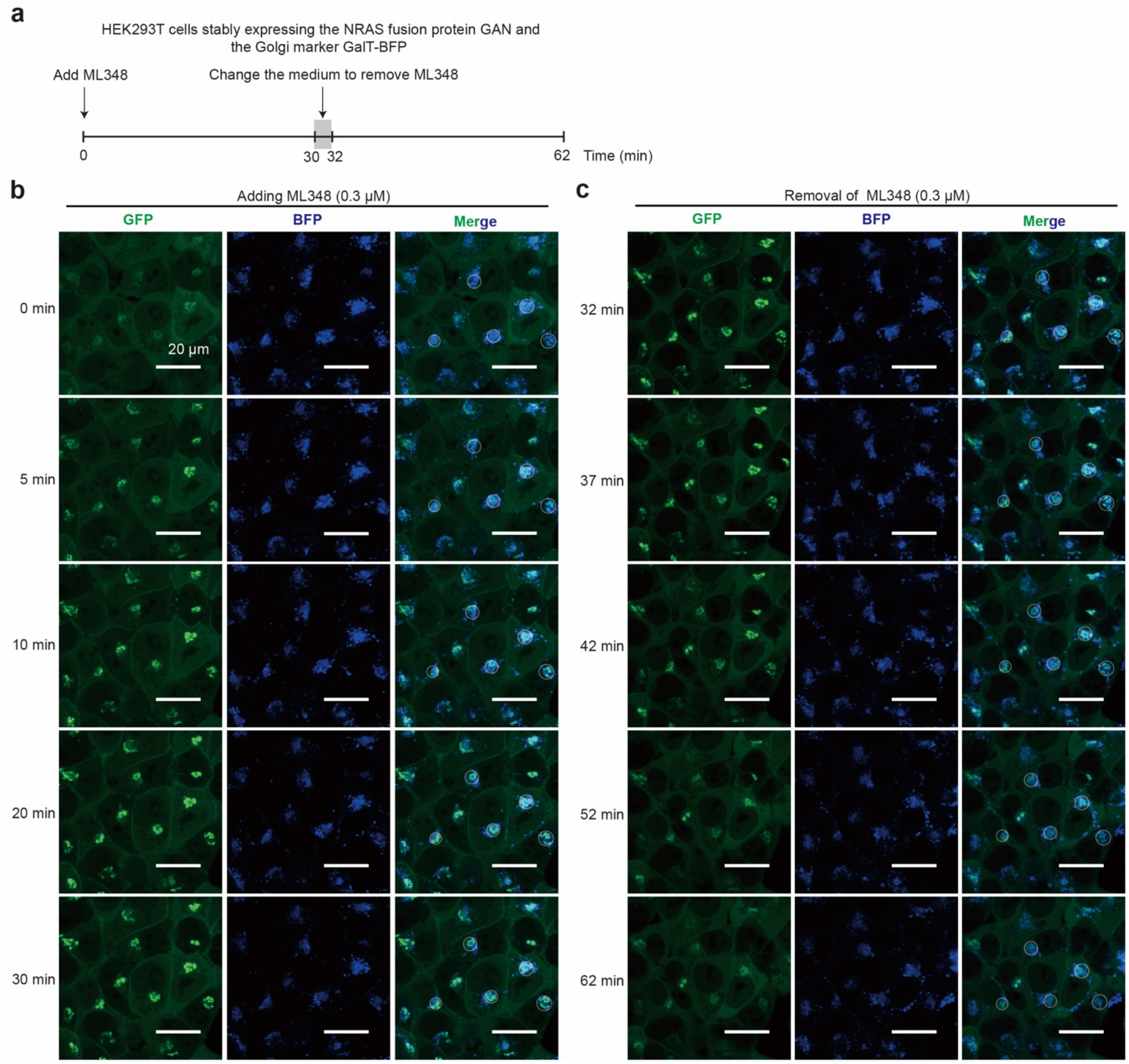


**Figure S5. ML348-induced re-localization of NRAS fusion protein GAN was reversed after removal of ML348 from the cell medium. a**, Illustration of the cell imaging process after adding or removing ML348 (same as Fig. 4a). **b**,**c**, Changes in the subcellular localization of GAN stably expressed in HEK293T cells after adding 0.3 μM of ML348 (**b**) and subsequently removing ML348 by changing the cell medium (**c**). The changes in the GFP signals in the regions indicated by white circles were quantified in Fig. 4d.

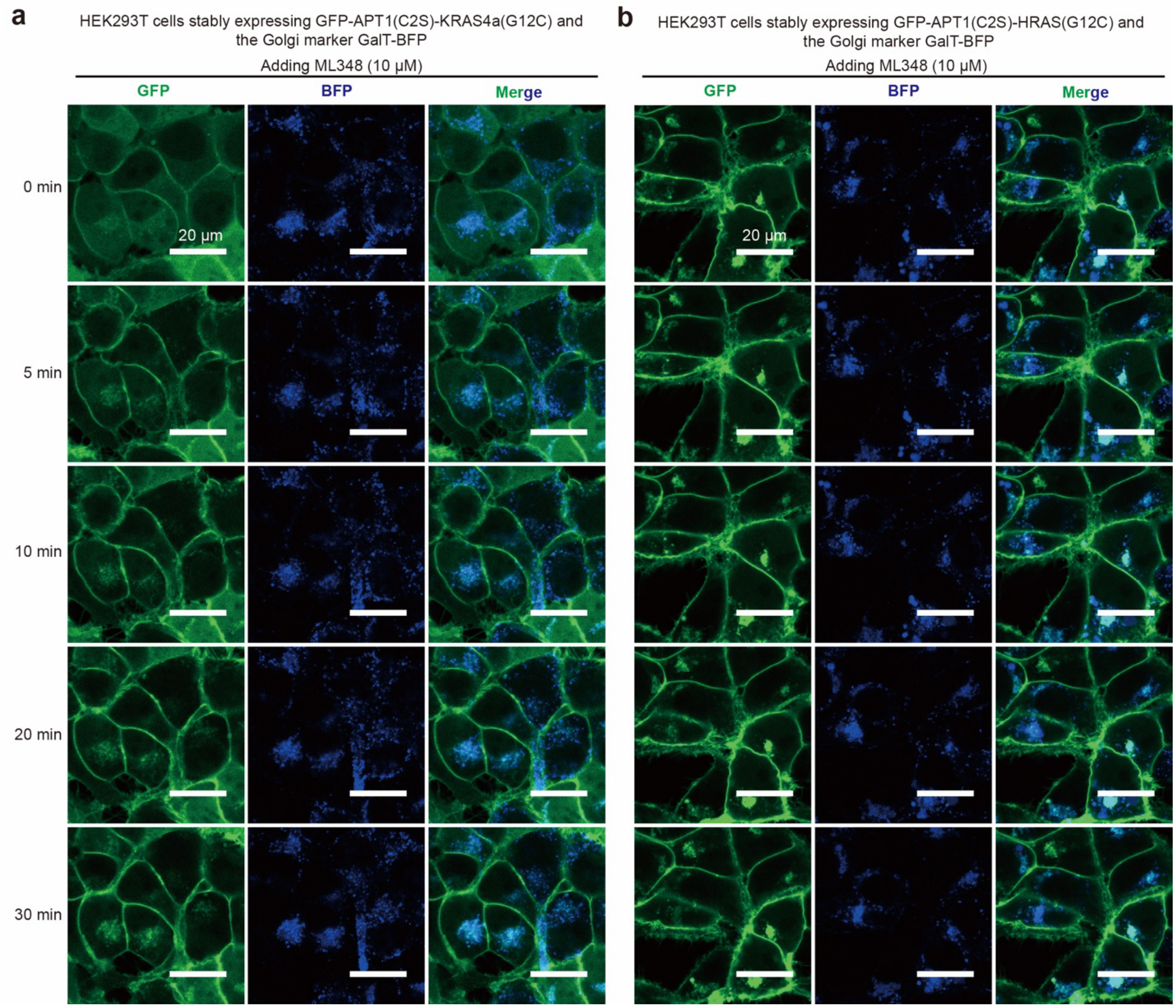


**Figure S6. Effect of fusing APT1 on the subcellular localization of the G12C mutants of KRAS4a and HRAS**. **a**,**b**, The KRAS4a(G12C) (**a**) and HRAS(G12C) (**b**) each with a GFP-tagged APT1(C2S) fused to the N-terminus were stably expressed in HEK293T cells. The cells also stably expressed the Golgi marker GalT-BFP. After adding 10 μM of ML348, the cells were imaged using ZEISS LSM 980 Confocal Microscope.

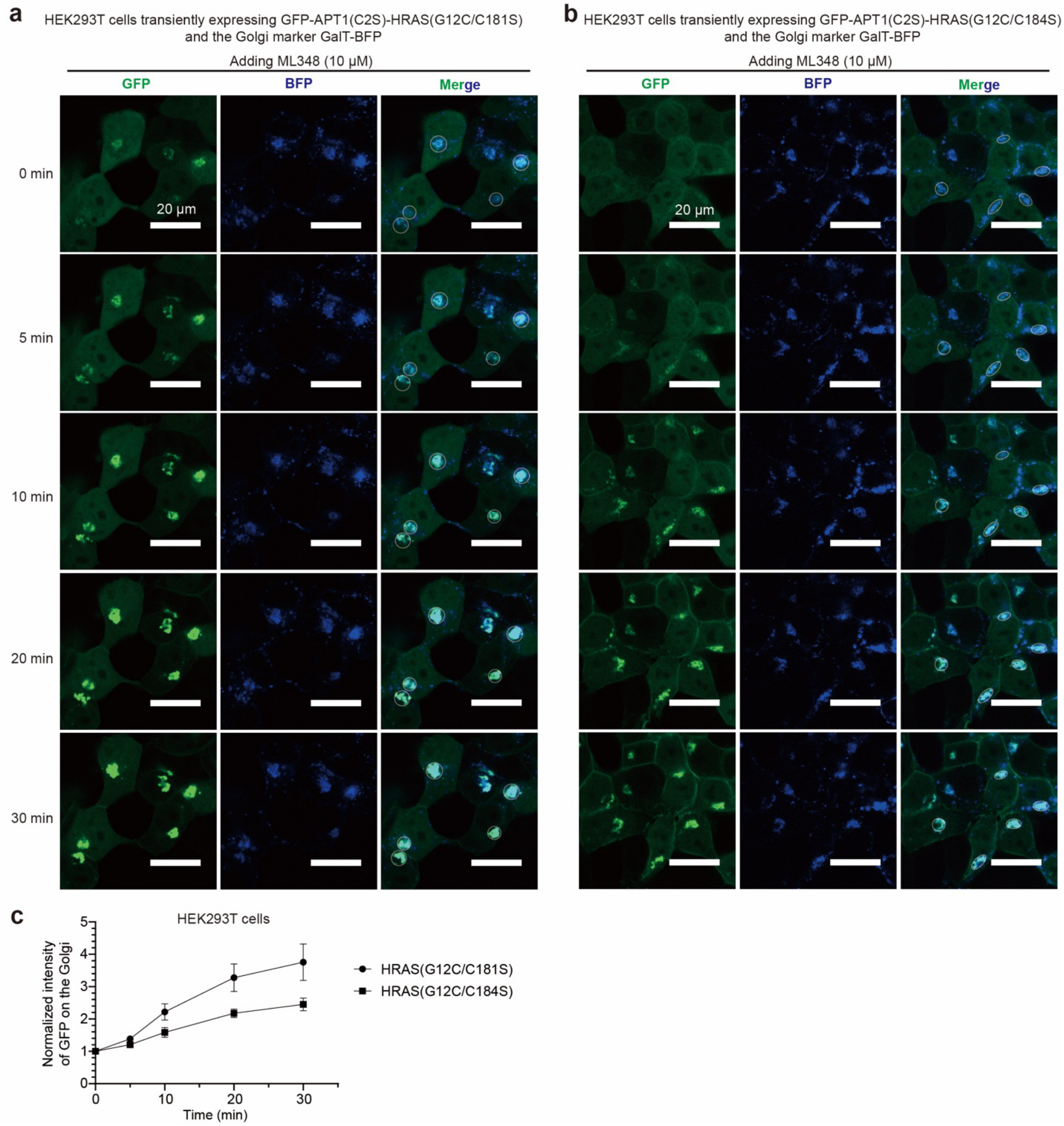


**Figure S7. Mutation of either C181 or C184 of HRAS(G12C) sensitizes it to APT1-induced depalmitoylation. a**,**b**, HRAS(G12C/C181S) (**a**) and HRAS(G12C/C184S) (**b**) each with a GFP-tagged APT1(C2S) fused to the N-terminus were transiently expressed in HEK293T cells. The cells also transiently expressed the Golgi marker GalT-BFP. After adding 10 μM of ML348, the cells were imaged using ZEISS LSM 980 Confocal Microscope. **c**, Quantification of the changes in the GFP signal on the Golgi in the HEK293T cells after adding 10 μM of ML348. The data in (**c**) represent the mean ± SEM of signals of five regions indicated by white circles in (**a**) and (**b**).
